# MIPMLP – Microbiome Preprocessing Machine Learning Pipeline

**DOI:** 10.1101/2020.11.24.397174

**Authors:** Yoel Y Jasner, Anna Belogolovski, Meirav Ben-Itzhak, Omry Koren, Yoram Louzoun

## Abstract

16S sequencing results are often used for Machine Learning (ML) tasks. 16S gene sequences are represented as feature counts, which are associated with taxonomic representation. Raw feature counts may not be the optimal representation for ML. We checked multiple preprocessing steps and tested the optimal combination for 16S sequencing-based classification tasks. We computed the contribution of each step to the accuracy as measured by the Area Under Curve (AUC) of the classification. We show that the log of the feature counts is much more informative than the relative counts. We further show that merging features associated with the same taxonomy at a given level, through a dimension reduction step for each group of bacteria improves the AUC. Finally, we show that z-scoring has a very limited effect on the results. These preprocessing steps are integrated into the MIPMLP - Microbiome Preprocessing Machine Learning Pipeline, which is available as a stand alone version at https://github.com/louzounlab/microbiome/tree/master/Preprocess or as a service at http://mip-mlp.math.biu.ac.il/Home

**Importance:** Microbiome composition has been proposed as a biomarker (mic-marker) for multiple diseases. However, a clear analysis of the optimal way to represent the gene sequence counts is still lacking.

We propose a simple and straight forward method that significantly improves the accuracy of mic-marker studies.

This method can be of use to merge two of the most important advances in biology in the last decade: Microbiome analysis, and the introduction of machine learning methods to biological studies.

## Background

Recent studies of 16S rRNA gene-sequences through next-generation sequencing have revolutionized our understanding of the microbial community composition and structure. 16S rRNA gene sequences are often clustered into Operational Taxonomic Units (OTUs) in QIIME I or features in QIIME II, based on sequence similarities. An OTU/feature is an operational definition used to classify groups of closely related sequences. However, the term OTU/feature is also used in a different context and refers to clusters of (uncultivated or unknown) organisms, grouped by DNA sequence similarity of a specific taxonomic marker gene[1]. In other words, OTUs are pragmatic proxies for “species” (microbial or metazoan) at different taxonomic levels, in the absence of traditional systems of biological classification as are available for macroscopic organisms. Although OTUs can be calculated differently when using different algorithms or thresholds, Schmidt et al. recently demonstrated that microbial OTUs were generally ecologically consistent across habitats and several OTU clustering approaches[2]. The same holds for the features in QIIME II.

OTU picking is the assignment of observed gene sequences to operational taxonomic units, based on the similarity between them and the reference gene sequences. The similarity percentage is user-defined (97% in QIIME I[3] and 99% in QIIME II[4]). This process has been an important step in the common pipeline for microbiome analysis. However, it may cluster components with different behaviors into the same unit, hiding component-specific patterns. There are many algorithms for OTU clustering such as: SortMeRNA[5], SUMACLUST[6], and swarm[7]. In this paper, QIIME I was used for OTU picking, giving a hierarchical representation for OTUs. However, the results are similar for QIIME II.

An important use of Microbiome samples is the development of Microbiome based Biomarkers (Mic-Markers), using Machine Learning (ML) tools. An important limitation of using bacterial features in machine learning is the feature hierarchy. The feature hierarchy is difficult to process and analyze due to the sparsity of the feature table (i.e. high number of the bacteria with 0 values in any typical samples). Moreover, even the limited number of observed features may be inflated due to errors in DNA sequencing[8].

Many applications of supervised learning methods, in particular Random Forests (RF), Support Vector Machines (SVM), Neural networks, and Boosting, and have been applied successfully to a large set of microbiota classification problems[9, 10, 11, 12, 13, 14, 15, 16]. However, little attention has been devoted to the proper way to integrate information from different hierarchical levels.

In different domains, Feature selection can be done by filtering methods, wrapper methods, or embedding methods[12]. Recent work on microbiota/metagenome classification, such as Fizzy[17] and MetAML[18], utilize standard feature selection algorithms, not capitalizing on the evolutionary relationship and the resulting hierarchical structure of features.

Fizzy implements multiple standard Information-theoretic subset selection methods (e.g. JMI, MIM, and mRMR from the FEAST C library), NPFS, and Lasso. MetAML performs microbiota or full metagenomic classification, which incorporates embedded feature selection methods, including Lasso and ENet, with RF and SVM classifiers.

These more generic approaches can be improved using methods explicitly incorporating the details of the taxonomy, such as Hierarchy Features Engineer (HFE)[19]. HFE uses all the taxonomy level and discards redundant features based on correlation and Information Gain (IG).

The goal of HFE was to formalize feature selection by systematically and reproducibly searching a suitable hypothesis space. Given a hierarchy of taxonomies, represented as a network where lower taxonomic (less detailed taxonomy) levels point to all the higher-level features belonging to the same lower taxonomic level, HFE is composed of 4 phases. 1) Consider the relative abundances of higher-order taxonomic units *i_k_* as potential features by summing up the relative abundances of their features they point to in a bottom-up tree traversal. 2) For each parent-child (low to high taxonomy) pair in the hierarchy, the Pearson correlation coefficient *ρ* is calculated between the parent and child feature frequencies over all samples. If *ρ* is greater than a predefined threshold of θ, the child node is discarded. Otherwise, the child node is kept as part of the hierarchy, 3) Based on the nodes retained from the previous phase, all paths are constructed from the leaves to the root (i.e., each feature’s lineage). For each path, the IG of each node on the path is calculated with respect to the labels/classes L. Then the average IG is calculated and used as a threshold to discard any node with a lower IG score or an IG score of zero. 4) The fourth phase deals with incomplete paths. In this phase, any leaf with an IG score less than the global average IG score of the remaining nodes from the third phase or an IG score of zero is removed. HFE is currently the algorithms incorporating the most information on the hierarchy, but it is a complex and computationally expensive approach.

Beyond the hierarchy, the ratio of sample number to feature number is of importance. In ML classification problems in general and in feature-based classifications that involve learning a “state-of-nature” from a finite number of data samples in a high-dimensional feature space, with each feature having a range of possible values, typically a large number of training samples is required to ensure that there are several samples with each combination of non-zero values[20]. This is typically not the case in feature-based ML.

A related, yet different issue is the input distribution. Many ML methods prefer a Gaussian distribution of input features. However, in features, a significant number of features have 0 values in many samples.

To address all these issues, We propose a general pre-processing algorithm for 16S rRNA gene sequencing-based machine learning, named MIPMLP (Microbiome Preprocessing Machine Learning Pipeline). The design principles of MIPMLP are:

- Optimization of machine learning precision, as measured here by Area under Curve (AUC) of binary classifiers.
- Ease of implementation.
- Explicit incorporation of detailed taxonomy in the analysis.
- Minimization of the number of tunable free parameters to avoid over-fitting to a specific dataset/task.

MIPMLP deals with the curse of dimensionality, skewed distribution, and feature frequency normalization. Different ML methods may obtain better accuracy with other feature selection pipeline. The advantage of this pipeline is that it can be used easily and it is not relying on labels like the HFE method[19]. Another advantage is that this pipeline can be used for every hierarchical feature representation task.

MIPMLP is available through an open GIT at https://github.com/louzounlab/microbiome/tree/master/Preprocess, and through a server at http://mip-mlp.math.biu.ac.il/Home

## Methods

MIPMLP proposes a pre-processing method based on three steps. Each step consists a choice from multiple options (Figure 1). The first step is the taxonomy level used for the representation and the taxonomy grouping method for the grouping step. The second step is a standardization step where one first decides if a log scale is performed or a relative normalization. The last step is a dimensions reduction step that specifies if a dimension reduction is performed, and if so which of PCA and ICA is performed.

**Figure 1:**
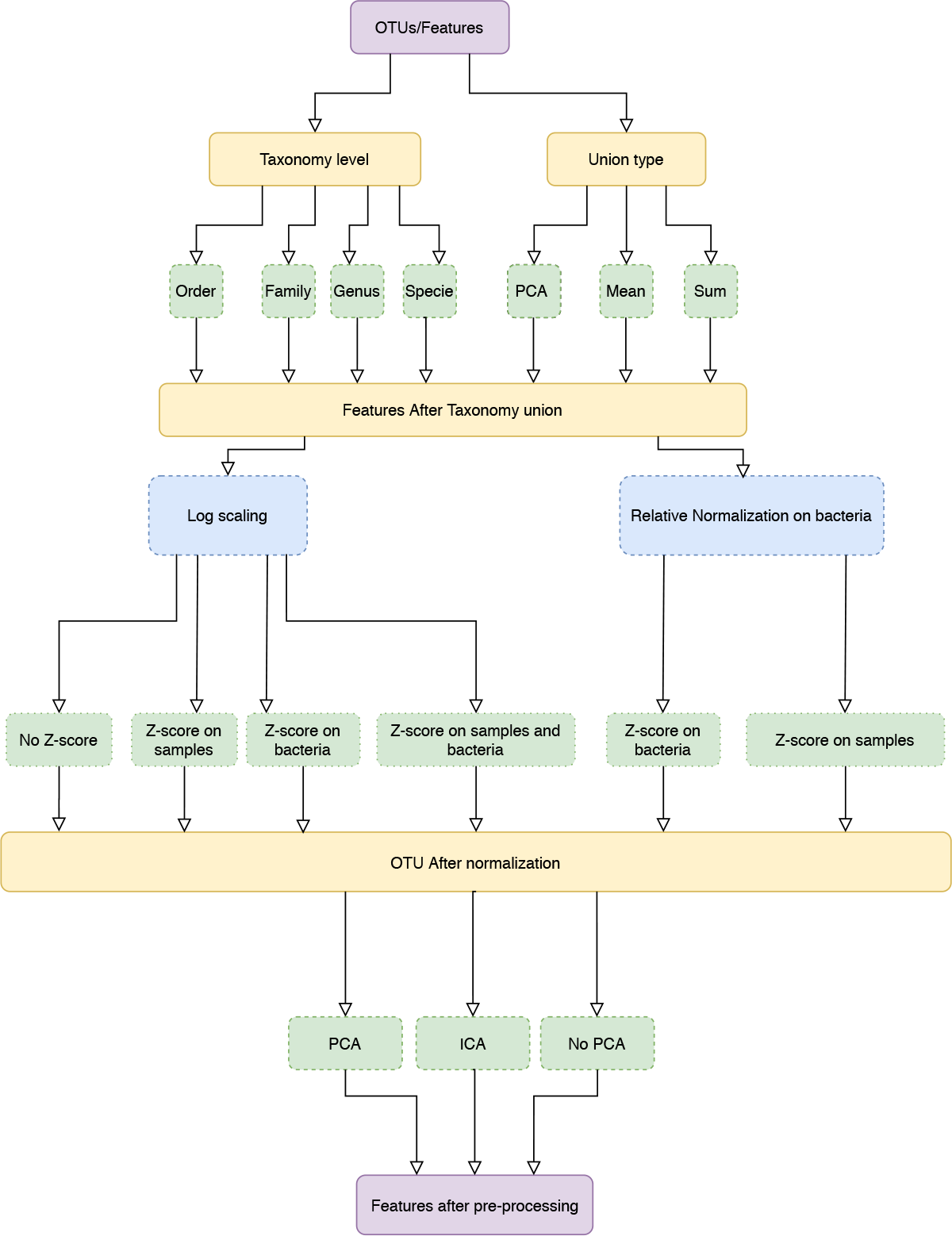
pipeline process diagram. The input is an OTU/Feature table and the appropriate taxonomy. The features are merged to a given taxonomic level. We tested three possible merging methods: Sum, Average and a PCA on each sub-group of features. Following the merging, we performed either a log scaling or a relative scaling. Following scaling, we performed z scoring on either bacteria or samples or both, and finally, we tested whether performing a dimension reduction on the resulting merged and normalized features improves the accuracy of predictions.

### Data sets

#### Il1 α

We investigated the connection between *IL1α* expression, microbiota composition, and clinical outcomes of induced colitis by using wild type and *IL1α* deficient [14].

#### Mucositis

In a collaboration with Sheba Medical Center, we serially collected 625 saliva samples from 184 adult allogeneic hematopoietic stem cell transplantation recipients and found microbial and metabolic signature associated with oral mucositis at different time points before and after the transplantation[21].

#### Progesterone

We demonstrated the dramatic shift in the gut microbial composition of women and mice during late pregnancy, including an increase in the relative abundance of Bifidobacterium. We showed the direct effect of progesterone elevation during late pregnancy on increased levels of Bifidobacteria[15].

### Taxonomy grouping

MIPMLP uses a taxonomy sensitive dimension reduction by grouping the bacteria at a given taxonomy level. All features with a given representation at a given taxonomy level were grouped and merged using different methods. We used three different taxonomy levels to group by (Order, Family, Genus). Three methods were used for the grouping stage:

1. Average of all the grouped features.
2. Sum of all the grouped features.
3. Merged using PCA following normalization, with a PCA on each group. Basically, all samples belonging to the same group are projected independently. We then use the projection on the PCs with the highest variance, explaining at least half the variance as the representation of the group, using the following algorithm:

**Algorithm 1.**
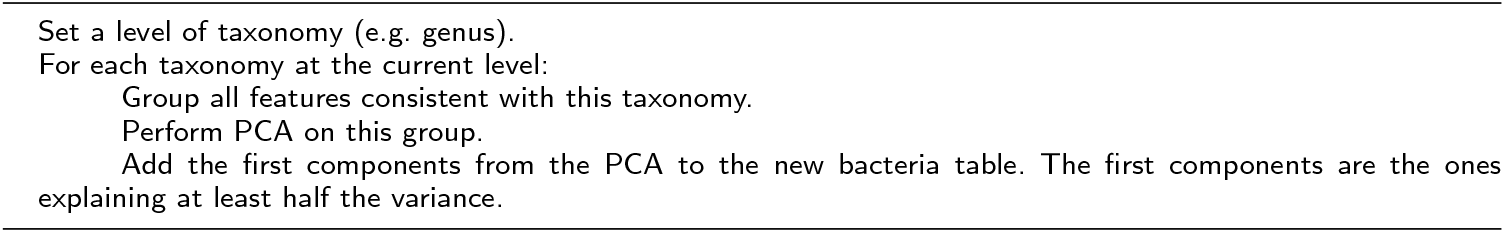
sub-PCA

0.1 Normalization and Standardization

Following grouping by any of the algorithms above, we tested two different distribution normalization methods. The first approach was to log (10 base) scale the features element-wise

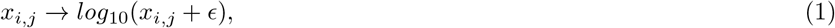

where *ϵ* is a minimal value to prevent log of zero values. The second one was to normalize each bacteria through its relative frequency:

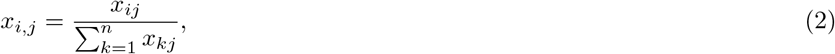

where *n* is the number of samples, *i* is the feature I.D. and *j* the sample I.D..

Following the log-normalization, we have tested four standardization possibilities: 1) No standardization, 2) Z-score each sample, 3) Z-score each bacteria, 4) Z-score each sample, and Z-score each bacteria (in this order).

When performing relative normalization, we either did not standardize the results or performed only a standardization on the bacteria (i.e. options 1 and 3 above).

A Z-score is defined as:

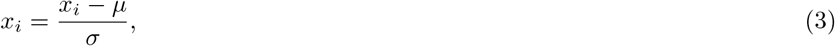

where *μ* is for the mean and *σ* is the standard deviation. (i.e. when applying a Z-score sample wise the mean is the mean of all the bacteria for one given sample. When applying Z-score bacteria wise the mean is the mean of all the samples for a given bacteria).

### Dimension reduction

PCA[22] and ICA[23] are dimensions reduction methods. While PCA is based on variance, ICA is based on Independence. After taxonomy grouping, normalization and standardization, we applied PCA, ICA, or none of them. The cut-off used for the accumulated variance was 0.7, and the same number of components were used for the ICA algorithm. In this paper Scikit-Learn.PCA and Scikit-Learn.FastICA were used as a coding framework[24].

### Machine Learning

To evaluate the different configuration, we used three different classifiers: 1) An SVM[25], linear classifier (with Scikit-Learn.svm.SVC) with a box-constraint of 0.1. 2) XGBOOST[26] with binary decision trees as the weak classifier with *n_estimators_* = 100, *γ* = 0.5 and *Minimum_childweight_* = 3, and 3) An Artificial Neural Network (ANN) [27] a feed forward network with two hidden layers. The first hidden layer of size 100. The second hidden layer had 100 neuron. The first activation function was ReLU[28] and the second activation function was a Sigmoid[29]. We used an Adam optimizer[30], with *Learningrate* = 0.005 and BCE (Binary Cross Entropy) loss function with *Batchsize* = 16.

The accuracy of all results was measured through the test set Area Under Curve (AUC) with ten-fold cross-validation. Note that the goal here is not to tune the hyperparameters of the learning, but rather to propose a pre-processing approach.

### Regression Model

To measure the contribution of each parameter of every step of the pipeline. For each classification method (SVM, XGBOOST, Neural Network), we regressed the test AUC on a one-hot representation of each specific pipeline configuration (unification level, method, normalization....) and calculated the AUC for the configuration. All the options were converted to a One-Hot vector representation so that each choice has a distinct coefficient. We then trained a multivariate linear regression on the train data set to predict the AUC for every configuration (i.e each row in the table stand for a different pre-process). The coefficients of the linear regression model were used to measure the contribution of each parameter to the pipeline. The linear regression can be described as:

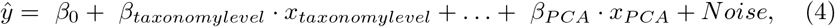

where *x_i_* is a binary input (0 if the parameter *i* was not used, 1 if the parameter *i* was used), *β_i_* is the coefficient of parameter *x_i_* (e.g. *x_taxonomy level five_* = 1 means that the union taxonomy level was the family level) and *ŷ* for predicted AUC.

Finally we subtracted the mean of the coefficients of every pre-processing step from each coefficient in the same step e.g.

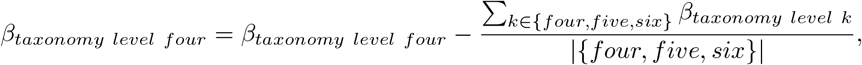

and used the *β_taxonomy level k_* as the contribution coefficient to the predicted AUC.

## Results

MIPMLP is a pipeline for 16S feature values pre-processing before machine learning can be used for classification tasks. To estimate the effect of different pre-processing steps coherently, we have studied multiple classification tasks. For each classification task, we used multiple machine learning algorithms. The algorithms’ hyperparameters were not tuned, since the goal was not to optimize the ML, but the pre-processing. As described in the method section, MIMLP contains four stages (Figure 1):

- Merging of similar features based on the taxonomy.
- Scaling the distribution.
- Standardization to z scores.
- Dimension reduction.

Following all these stages, a binary classification task was performed (see method section). We then computed for each data-set and each classification method the accuracy of the classification, through the AUC (Area Under ROC curve) the ROC curve is created by plotting the TPR (true positive rate - sensitivity) against the FPR (false positive rate - 1-specificity) at various threshold settings (Figure. 2). The average AUC for each task was computed using 5-fold cross-validation (i.e. split the data with test size of 20% and average 5 splits). The training/test division was fixed among classification tasks for each data-set. the first step of MIPMLP involves two choices, the first one is the taxonomy level and the second is taxonomy grouping type (Figure 1 third row):

1. The taxonomy levels used for merging are Order, Family, and Genus. We did not analyze at the species level, since this consistently gave worse results than lower levels (less detailed taxonomy).
2. The methods of merging. Three different methods were tested:

1. Sum of all features associated with a given bacteria (at the level chosen above).
2. Average of all features associated with a given bacteria, as above.
3. A more complex approach was to reproduce the variability in each level, by performing a dimension reduction on all samples and all features associated with a bacteria, and representing the bacteria by the projections reproducing almost half the variance. Note that this method was performed after scaling (to have a normal input distribution for the PCA). The second step involves two choices of scaling. The first one being relative scaling, which is currently the standard in most studies - the division of each feature frequency by the sum over all feature frequencies in the same sample. An alternative approach is scaling all feature frequencies to a logarithmic scale. Since a large number of features have 0 frequency in many samples. A minimal value (ϵ = 0.1) was added to each frequency.

**Figure 2:**
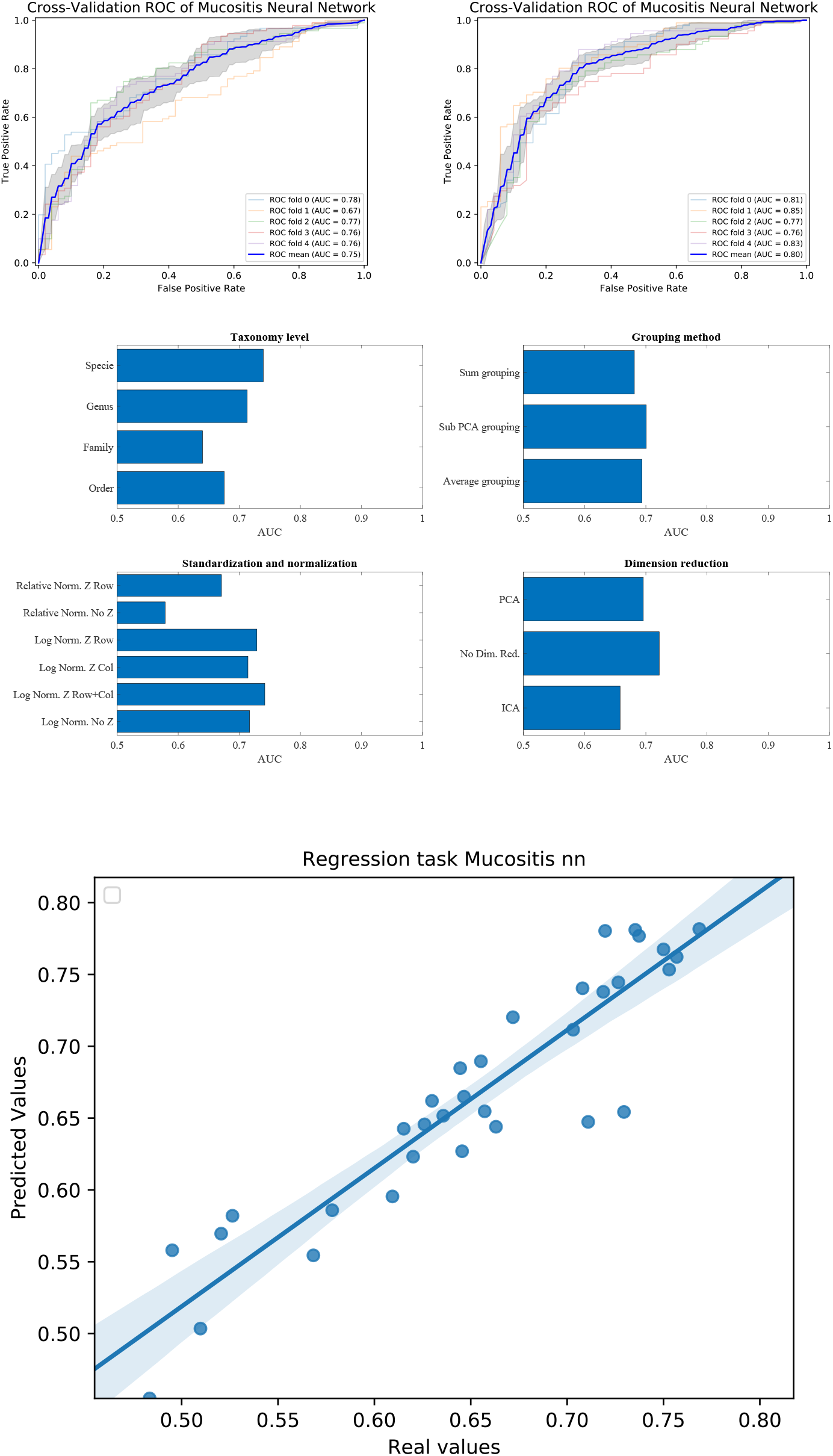
Upper plot typical ROC and the effect of preprocessing. The right plot is using the sub-PCA merging method, while the left plot us using the average merging method. The right upper plot has a higher AUC than the left one. Middle plots - average AUC defined as average AUC using one feature (e.g. one taxonomy level) and all other combinations (e.g. merging methods, normalization etc). Lower plot - Predicted AUC in linear regression vs real linear regression.

The third step involves normalization. There are two main arguments for normalization. To ensure that all features entering the machine learning have equal average and variance. This could be obtained by z-scoring each feature to zero average and unity variance. An alternative normalization would be to ensure that differences in the amount of genetic material would not affect the results. This would require a z-scoring over the samples. Finally one could propose doing both types of z-scoring. We have tested all three possibilities.

The last step of the analysis was to perform dimension reduction over the resulting projections. We tested three options: 1) No dimension reduction, 2) PCA, 3) ICA.

To exemplify how we test the combination of pre-processing steps, we follow an example in one dataset and one learning method.

### Example on Mucositis prediction prognosis from pre-transplant microbiome samples

Let us follow the analysis for an Artificial Neural Network (ANN) based prediction of the emergence of Mucositis following bone marrow transplant in leukemic patients[21]. We present the differences between the configuration in every preprocessing steps through their influence on the prediction precision. We focus in this section on the Mucositis prediction and ANN. Similar results were achieved using other data sets and classifiers, as further detailed below.

To test that the pre-processing indeed affects the test-set AUC value, we tested all possible combinations of all pre-processing steps. For each combination, we averaged the AUC over all training/test divisions (Figure 2. upper plots). We then evaluated the AUC obtained using a specific value in a given step, and all options in all other steps (Figure 2. Middle plots). For example, to estimate the expected accuracy when using a genus-level representation, we averaged the AUC Of all evaluations using a genus-level representation, and all possibilities on all other choices. One can see that for the ANN and the Mucositis prediction that making a taxonomy grouping of sub-PCA (the novel method presented here) and a genus based representation gives the optimal AUC. Using family or Order taxonomy levels decreases the AUC by 0.03-0.08, and using other methods than sub-PCA decrease the AUC by 0.05. Similarly, using a logarithmic normalization and z scoring both columns and rows is the optimal approach for this dataset on average. However, the effect of Z-scoring is minimal. One can further see that any dimension reduction reduces the test set accuracy, but there is no clear difference between ICA and PCA. These results are similar to the results obtained using more rigorous methods as follows.

To address the effect of combinations of pre-processing steps and their inter-dependence, we used a regression-based approach, where the contribution of each step to the test set accuracy is represented by its coefficient in the regression. This is performed by representing the set of options via a one-hot representation, and performing a linear regression of the test set average AUC (averaged over cross-validations) over the one-hot vectors. The resulting coefficients are not unique (since the one-hot representation matrix is not full rank). Thus, to compute the relative effect, we normalized the average effect of the interchangeable coefficient to 0 (for example the representation level, or the dimension reduction). To test that such a correlation can give meaningful results, we computed the expected and observed test AUC in the ANN Mucositis prediction (Figure 2 lower plot). Indeed the real AUCs are tightly correlated. The resulting coefficients can be used to assess the effect of a pre-processing step. We then applied the same method to all data-sets and all learning methods.

### Comparison of different data-sets and learning methods

To test the effect of the pre-processing in general and with different ML frameworks, We computed the regression coefficients for each data set and each ML methods. We present the distribution of coefficients of every data set in a box-plot, for each classifier separately (Figures 3,4,5).

**Figure 3:**
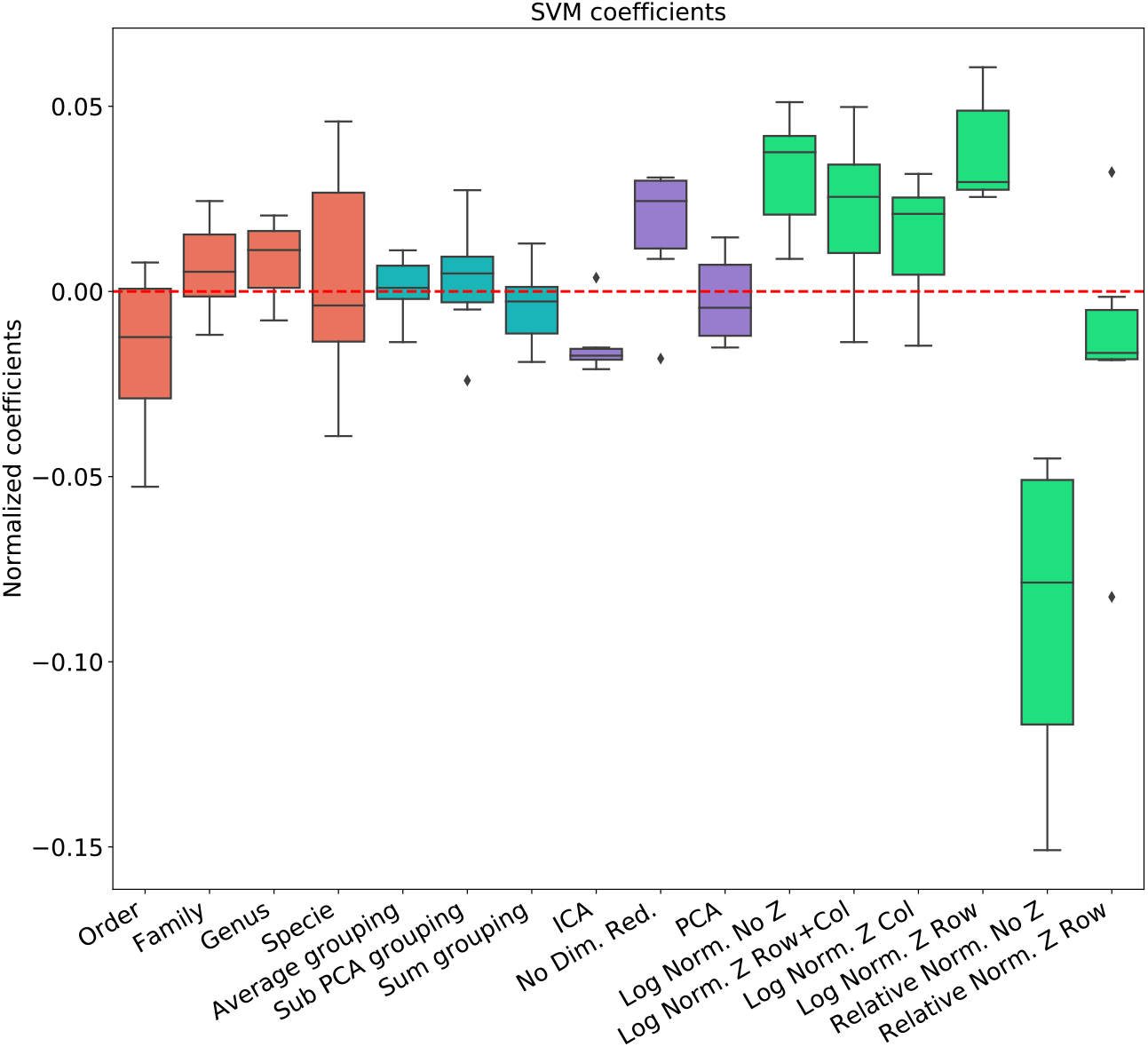
linear regression coefficient for SVM classifier. Coefficients are the contribution of a choice to the total AUC. Each group of coefficients is marked by a different color and normalized to 0. The following two figures follow this figure, but for different classifiers. The regression is over all parameter combinations, including the choice of taxonomy level (red), the grouping method (blue), the dimension reduction method (purple) and the normalization method (green). Since not all normalization and standardization methods are possible, we opened all tested combinations.

**Figure 4:**
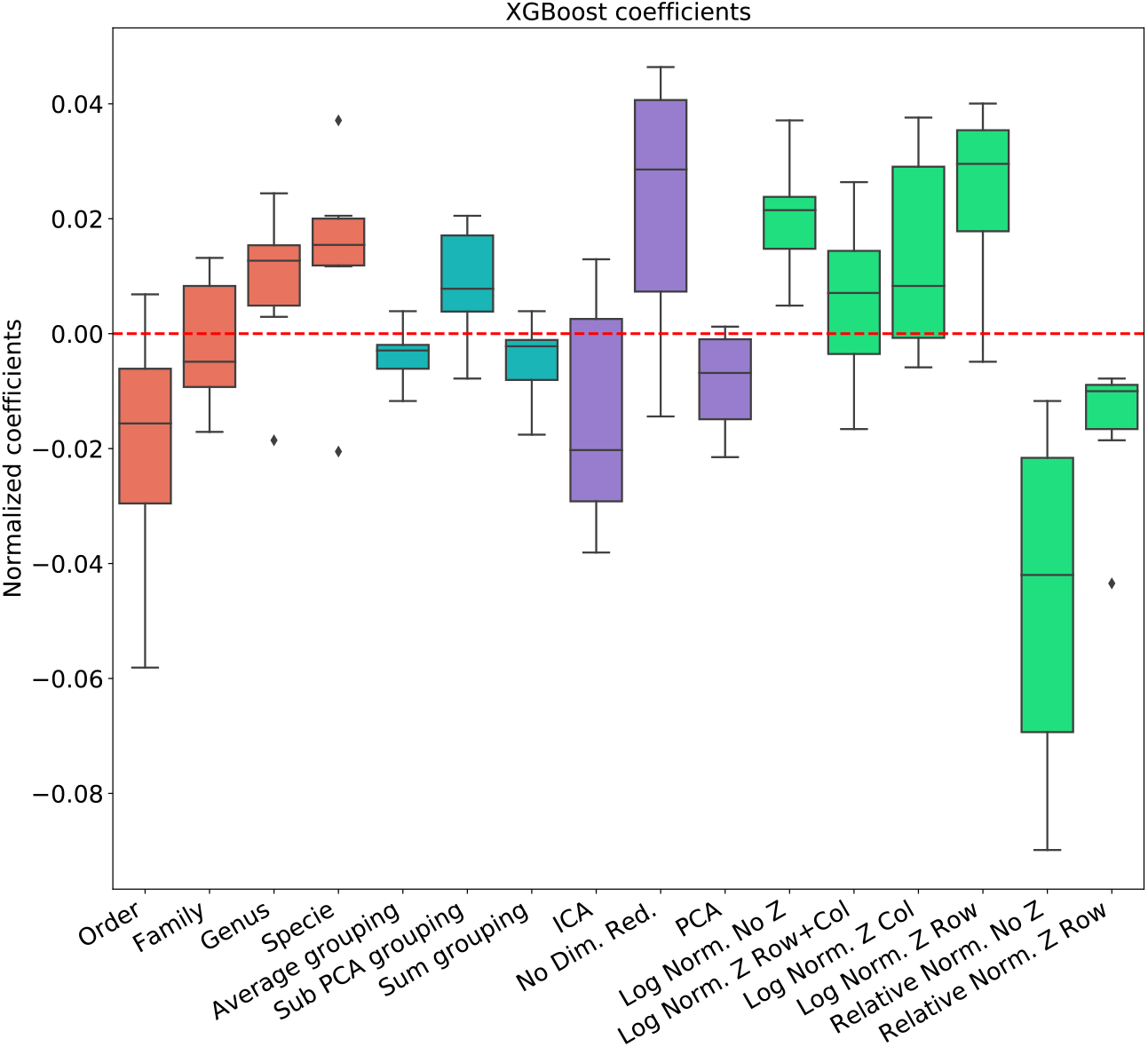
linear regression coefficient for XGBosot classifier.

**Figure 5:**
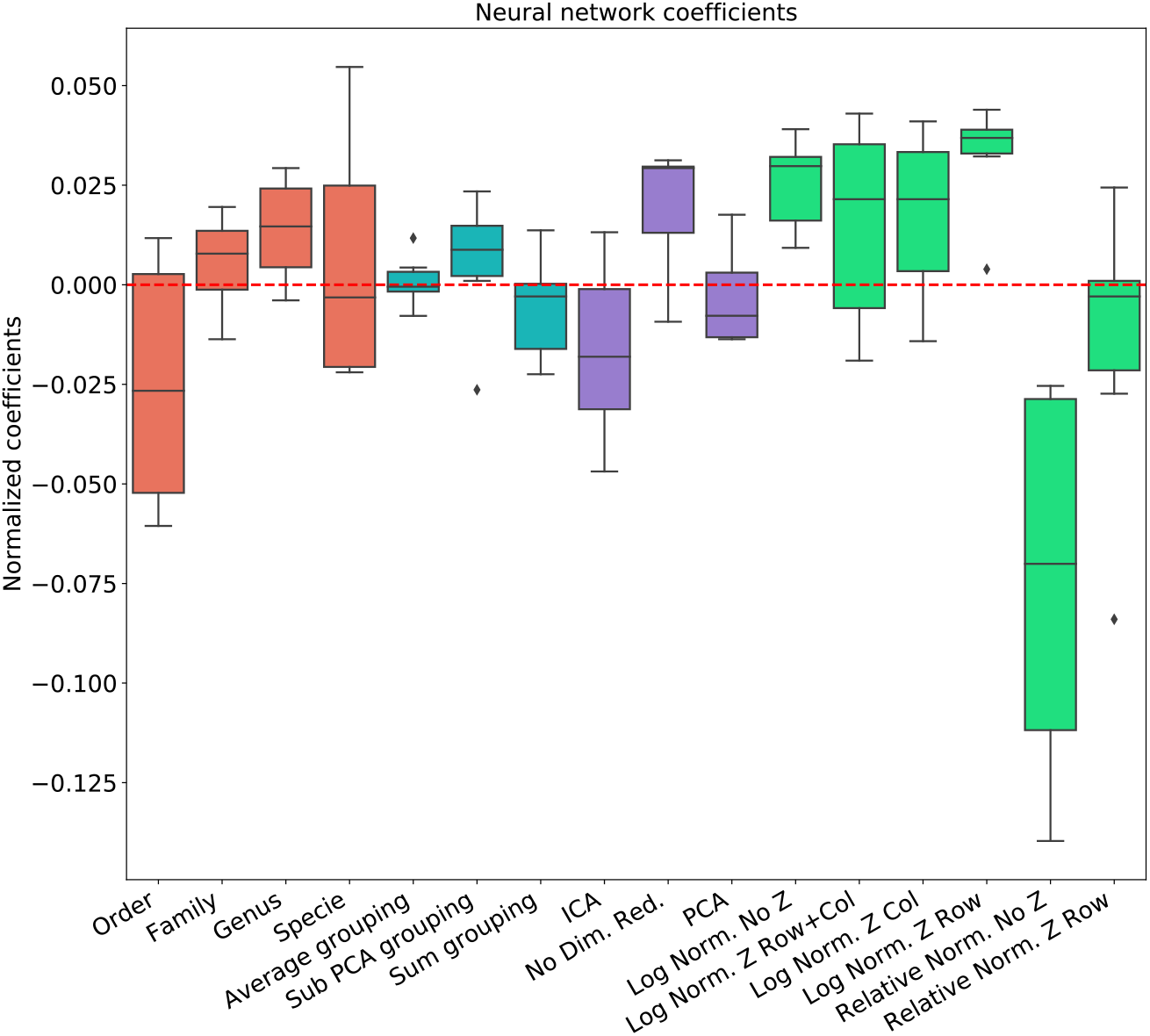
linear regression coefficient for MLP classifier.

The results are highly consistent among the different learning methods, with the following conclusions:

- The main effect is the effect of normalization. Relative normalization reduces the AUC on average by more than 0.05 compared with log scaling the data.
- A Genus taxonomy level representation is typically the best, and reducing to lower orders can further reduce AUC by 0.02-0.03.
- All dimension reduction algorithms reduce the AUC by around 0.02.
- The sub-PCA method to merge features increases the AUC by 0.01-0.02 compared with the sum or the average.
- Z-Scoring has a minimal effect, and the precise Z-scoring performed is of limited importance.

We thus suggest that OTU data should be pre-processed at the Genus taxonomy level with log scaling, using the sub-PCA algorithm (presented in the method section), and no further dimension reduction. Note that the cumulative addition of approximately 0.1 by this combination may be crucial for many ML applications.

Another much more complex algorithm for microbiome ML pre-processing is the HFE algorithm that was suggested in [19]. To compare MIPMLP and HFE, we used an SVM classifier with a box constraint of 0.01 and a linear kernel, we tested the results on 5 different data sets each of them was split with 7-fold cross-validation.

As can be seen in figure 6, HFE tends to over-fit very easily. we can assume that the reason for that is that HFE is based on train labels for computing thresholds and correlation at the pre-processing step, before the learning methods. Also, in general, for MIPMLP and HFE the AUC values are relatively close, but MIMLP is much simpler and computationally effective.

**Figure 6:**
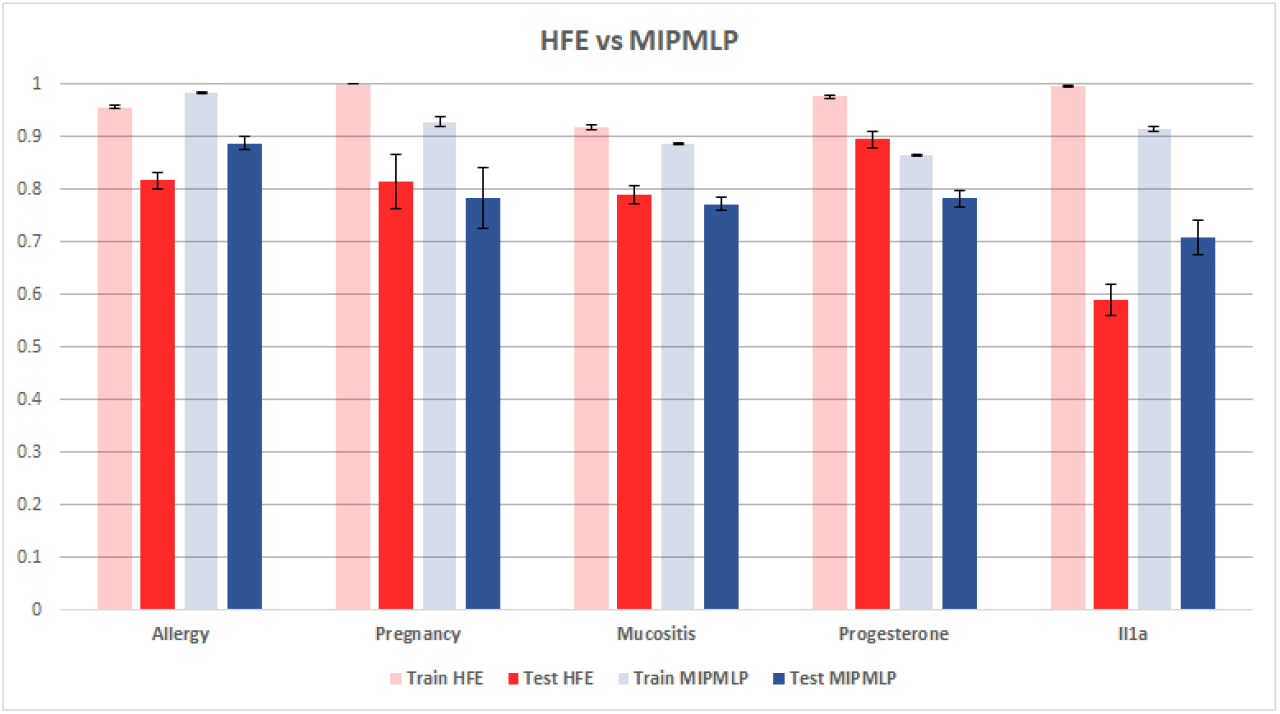
HFE and MIPMLP mean AUC with standard errors bar. shaded bars are training set and full bars are test set. Error bars are standard errors. The y axis is AUC. Different groups of bars are different datasets.

## Conclusion

We have presented here MIPMLP a computationally efficient framework to preprocess 16S feature values for ML based classification tasks. While MIPMLP allows for multiple choices at each stage of the pre-processing, a consensus method emerges which typically gives the optimal AUC, which is:

1. Use a Genus level representation.
2. Perform a log-transform of the samples.
3. Merge all features belonging to the same genus through a PCA on these specific features.
4. Do not perform other dimension reductions.
5. Z-scoring has a minimal effect if any on the results.

The importance of the log-transform suggests that the information of the most abundant species is limited. Instead, the relative change in the frequency of all species, even rare ones should be used. The genus-level presentation suggests that the prediction is not based on any specific bacteria, but rather on some more general aspects, such as the metabolite usage and production, or the association with inflammation[21]. Finally, the need for PCA instead of average/sum when merging a genus, suggests that treating all features are equal is sub-optimal. Instead, features contributing to the variance between samples should receive more importance.

## Discussion

The microbiome is now widely used as a biomarker (mic-marker) in the context of ML-based classification tasks. However, very limited attention was given to the optimal representation of 16S based features for such classification tasks. While features are used as a representation of species, in reality, a feature is an abstract representation of bacteria clustered based on the similarity of one protein. As such, we propose that a higher level of representation would give a higher accuracy in ML tasks. We then propose a formalism to integrate feature expression levels for such a representation.

Similarly, the expression level in sequencing experiments is not a direct measure of the number of bacteria, but instead the result of multiple experimental and computational stages, including the extraction of the genetic material, primer specific PCR amplification[31], and computational sequence quality control. Our results suggest that comparing the absolute feature frequencies (or relative frequencies) in different experiments leads to lower accuracy than measuring the fold change, as expressed by differences in the logged frequencies. Indeed, in most of our recent results [14, 16, 21, 32], we found that such a log normalization is essential.

As many non-computational scientists are now entering the field of ML in microbiome studies, we believe that MIPMLP will help standardize the use of ML in microbiome studies. We feel that the processing steps described throughout the manuscript will allow for better prognosis and diagnosis as they focus on the common features in the microbiome at different taxonomic levels. Employing such an approach as we described here will allow moving microbiome-based prediction of disease states from bench to bedside.

## Author’s contributions

YJ and YL wrote the manuscript and the code, as well as performed part of the analysis.... AB and MB developed some of the methods.... OK wrote a part of the manuscript and performed the experimental analysis.

## Acknowledgements

OK and YL acknowledge the Food IoT grant and the Bar Ilan DSI grant…

